# Development of an open source program to identify the size of objects using bright-field microscopy - an automated calibration tool

**DOI:** 10.1101/2021.04.30.442178

**Authors:** Gerard Glowacki, Alexis Gkantiragas, Brooke Brett-Holt, Daniel Mihalik, Peter He

## Abstract

In light microscopy, eyepiece graticules are commonly used to gauge the size of objects at the micron scale. While this is a relatively simple tool to use, not all microscopes possess this feature. Furthermore, calibrating an eyepiece graticule with a stage micrometer can be time-consuming, particularly for inexperienced microscopists. Similarly, calculating the size of individual objects may also take some time. We present an open-source program to determine the size of objects under a microscope using Python and OpenCV. Taking photos of a stage micrometer under a microscope, we identify gradations on the micrometer and calculate the distance between lines on the micrometer in pixels. From this, we can infer the size of objects from bright-field microscopy images. We believe this will improve access to quantitative microscopy techniques and increase the speed at which samples may be analyzed by light microscopy. Future studies may aim to integrate this with machine learning for object identification.

## Introduction

Accurately calculating the size of objects under a microscope is important for an enormous range of applications, from measuring intracellular structures to particulate analysis in forensic science [1] [2]. In our work, the precise measuring of diameter is an important factor in the classification of pollen grains [3].

Eyepiece graticules, in combination with stage micrometers, have been in use for decades in microscopy to provide a calibrated size scale [4]. Haemocytometers, first developed in 1902 [5] and similar devices can also be used to provide scaling for small objects such as cells. Perhaps the simplest form of calibration is one in which the field of view is compared with the total size of a stage micrometer. By estimating the size of the micrometer in relation to the field of view, one can (very approximately) guess the size of objects at that magnification. Alternatively, most microscopists measure the size of one eyepiece unit using a stage micrometer. Following this calibration one may calculate the size of objects by comparing them to the size of the micrometer. It is advisable to re-calibrate the microscope with each use.

While previous studies and tools use manual calibration or rely on microscopes with built-in calibration [6], this has limited scope for lower-end microscopes. A common calibration technique relies on the microscope stage moving by 1 micron in a given direction, and this is then translated into a particular distance on the eyepiece/field of view - thereby calibrating the microscope [7]. However, this technique has several limitations - namely that it assumes that the microscope has a movable stage and that this stage can be moved accurately to within a micron. Previous studies on automated calibration have also developed sophisticated calibration techniques for electron tomography [8, 9] and z-stack microscopes [10].

Our group works on automated detection and classification of pollen [11, 12]. In our previous studies, we did not take absolute scale into account and relied entirely on relative size. Therefore, this study aimed to develop a novel, robust tool to use microscopy images of a stage micrometer to calibrate a light microscope. While this tool was initially developed for pollen analysis, it is applicable to light microscopy more generally. In this paper, we call the real-world distance represented by a pixel the ‘pixel distance’ of an image.

## Materials and Methods

### Materials

The microscope used was an AmScope T360B LED Trinocular Biological Compound Microscope with 40X-2000X magnification. Images were taken using a Samsung A6 16MP rear camera or a Xiaomi A2 rear camera. We attached the smartphone to the microscope using a Solomark Universal Cell Phone Adapter Mount. The stage micrometers were purchased from MUHWA for both a standard micrometer and a micrometer with multiple scales.

### Statistical analysis

Confidence interval (CI) was calculated using the Student’s t-distribution. Confidence interval was set to 95%.

### Methods

Pictures were taken using a smartphone device attached to the microscope.

We identify the gradations on the stage micrometer by performing a canny edge detection followed by a cross shaped dilation kernel to close any holes in the edges [13] and filling the entire image, leaving areas within contiguous edges unfilled. We then fit rotated rectangles to these contours. We obtain the endpoints of these gradations using the midpoints of the shorter sides of these rectangles. We reject any rectangles without a sufficiently high aspect ratio. A ratio of 1:20 was used as a default (which worked well for our micrometer) but this parameter can be tuned in the software.

To calculate the distance between two gradations we select the midpoint of one gradation and projected this point onto the other gradation. We then calculate the distance between these two points (figure 1). Next, we compare every gradation to every other line and record the shortest distances. We reject a user-defined percentage of the longest and shortest distances, then average the remaining distances to find the average shortest distance between two lines. We divide the true distance between the gradations provided by the manufacturer by this average shortest distance to find the pixel distance.

**Figure 1.**
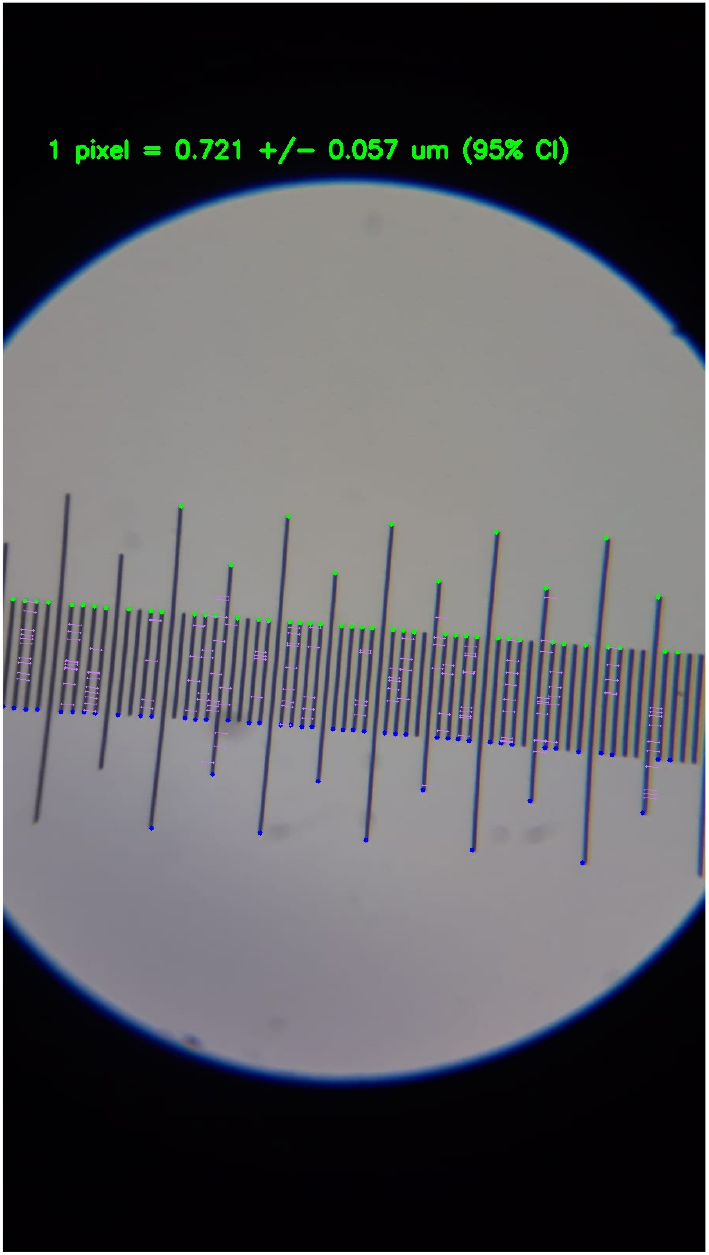
Distance pairs. Visualisation of pairs of distances between parallel lines following culling of exceptionally long and short distances

We validate these results by manually counting the pixel distances between a sample of lines on this image and the result determined by our algorithm. We conduct this using two different micrometers and smartphone cameras.

### Availability of data and materials

All data and code used are publicly available and have been uploaded to Git-Lab (https://gitlab.com/guunterr/microscope-scale-calibration). All materials used in the study are available via online retailers such as Amazon.

## Results

We tested our algorithm by manually counting pixels to calculate the pixel distance and comparing this distance to the value obtained by our algorithm. This test was repeated at 2 different magnifications (100x and 250x), using 2 different smartphone cameras and on 2 different stage micrometer slides: one where all the graticules are in a line (which we refer to as a *line* micrometer) and one where the graticules are arranged in a cross around a central grid (which we refer to as a *cross* micrometer). Results are shown in Table 1. It is worth noting that the 95% confidence interval always contained the ground truth value (figure 2).

**Table 1.** Results for automated and manual calibration using line and cross micrometers on 2 different microscopes. Values for automated calibration are given +/− the standard deviation.

| | Manual($\mu\text{m}/\text{pixel}$) | Algorithm ($\mu\text{m}/\text{pixel}$) | error |
| --- | --- | --- | --- |
| 100x, Samsung, Line | 2.0530 | REJECT | - |
| 100x, Samsung, Cross | 1.7492 | REJECT | - |
| 250x, Samsung, Line | 0.7919 | $0.7897 \pm 0.0723$ | 0.28% |
| 250x, Samsung, Cross | 0.7310 | $0.7319 \pm 0.0686$ | 0.12% |
| 100x, Xiaomi, Line | 0.5849 | $0.5850 \pm 0.0153$ | 0.01% |
| 100x, Xiaomi, Cross | 0.5743 | $0.5764 \pm 0.0213$ | 0.37% |
| 250x, Xiaomi, Line | 0.2368 | $0.2430 \pm 0.0072$ | 2.62% |
| 250x, Xiaomi, Cross | 0.3022 | $0.2967 \pm 0.0831$ | 1.82% |

**Figure 2.**
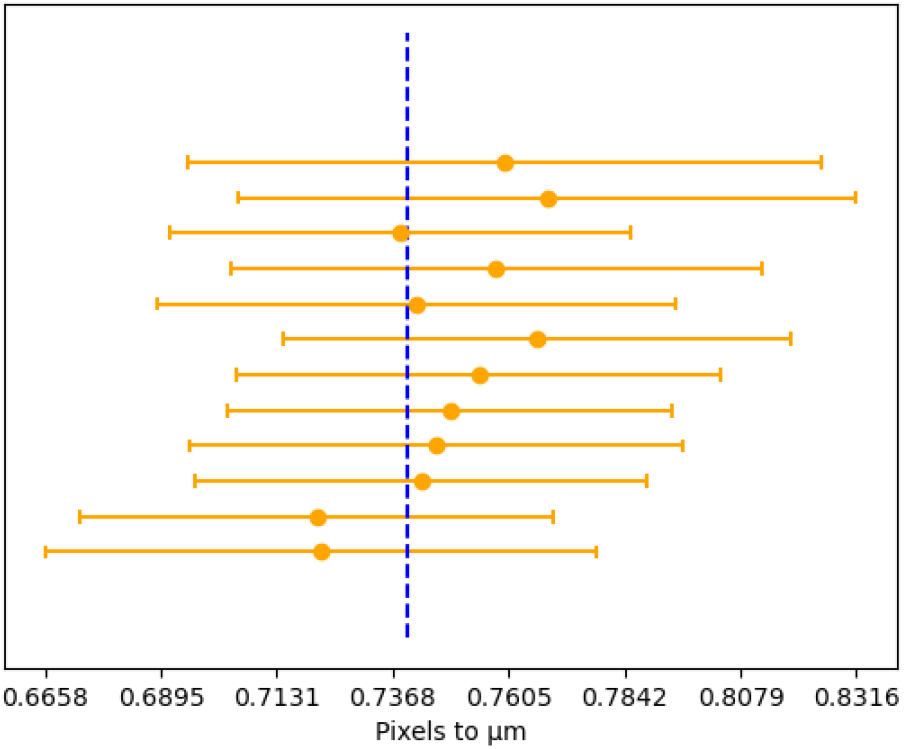
Confidence intervals. Visualisation of confidence intervals around means (orange) with respect to the true value (blue line) for a sample of images. Images of a line type micrometer were taken at 250x with a Samsung phone

Unsurprisingly, accuracy increased with increasing magnification since the distances between lines were represented by a greater number of pixels. Accuracy was not noticeably impaired by changing camera, though lower magnifications on the Samsung were rejected by the program as it was unable to resolve the edges on the microscope’s gradation, likely due to noise interfering with the edges. Accuracy was not noticeably affected by the use of different graticules as crossed and lined graticules yielded similar results. Slightly out of focus or noisy images did not significantly hinder the calibration, likely due to the Gaussian smoothing applied as part of the canny edge detector. Accuracy was not decreased even by significant artefacts such as uneven illumination, edited in dirt, writing, drawings and numbers on the slide (figure 3).

**Figure 3.**
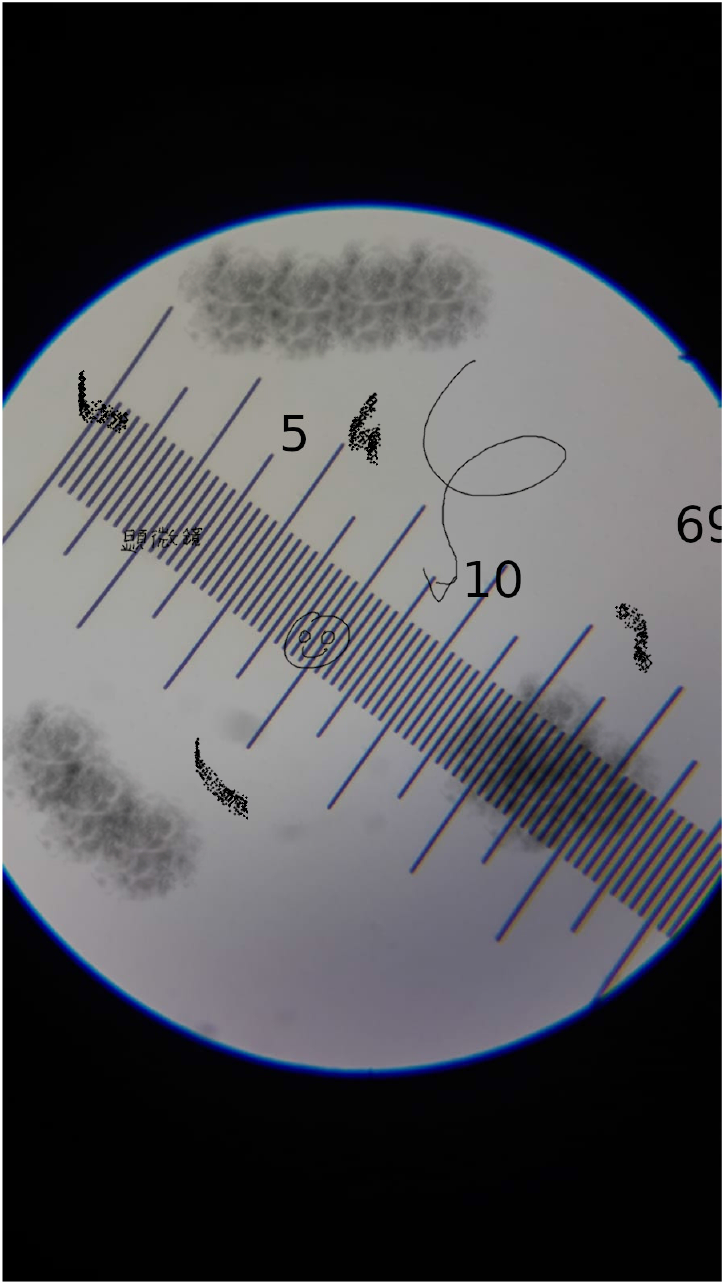
Robustness against small artefacts. Both images returned distances whose confidence intervals contained the respective ground truth

To enable easy use of our tool, we developed a user interface. Users are posed 2 questions. 1) What type of stage micrometer are you using? 2) What’s the distance between the lines? Finally, users are prompted to upload their images and from this pixel size can be calculated. Parameters including the expected aspect ratio of the rectangles fitted to gradations, the percentage of distances to reject and the size of the canny kernel can also be passed to the software through an options dialog. It is our hope that this sufficiently accounts for possible variations between micrometers. A video detailing the process is shown in supplementary video 1.

## Limitations

While we have tested our approach on two different cameras, there may be some devices that are incompatible with our approach. Furthermore, it is possible that different microscopes may encounter difficulties that we had not anticipated. Similarly, our tool is likely to be incompatible for other microscope types (e.g. phase contrast). However, it is likely to have applications in dissection microscopy - which regularly use micrometers to measure the size of objects such as insect limbs [14]. Our tool still requires the use of a stage micrometer which may not be available to all microscope users. However, it eliminates the need for using both a stage and eyepiece graticule reducing costs several-fold. On the other hand, the need to use a computer and camera (including smartphone cameras) may hinder some users.

While experiments with objects such as floating text and numbers overlaid on the images showed little change in results, comparison across different microscopes and lenses with aberrations as well as those with differing cleanliness are needed to assess the robustness of this tool. Finally, while we do eliminate the need for manual calibration, the actual object measurement still requires some input from the user. We hope that future studies may overcome these limitations.

While we could have employed a deep learning approach, we concluded that this would limit the accessibility of the tool to those with access to more powerful devices. Nonetheless, we cannot exclude the possibility that this would yield more accurate results.

Ultimately, use by specialists and non specialists will be required to optimise and validate our tool.

## Conclusion

We developed an accurate and novel tool to calibrate microscopes using only a stage micrometer. This has possible applications as a learning tool in classroom settings since measurement of hair, onion cells or other biological specimens are routine classroom experiments. Furthermore, we hope that it may eliminate variability in measurements caused by differences in individual users. Moreover, our tool will make absolute scale measurements in object identification driven by machine learning far easier since images taken will not be distorted by the presence of an eyepiece graticule. Future studies could try to integrate this with machine learning driven object identification. It is our hope to apply this tool to automate the classification of pollen using bright field microscopy.

## Author’s contributions

AG, BBH and PH conceived of the study. GG and PH conducted the experimental work. DM, PH and GG wrote the code. AG, PH and GG wrote the manuscript. All authors read and approved the final manuscript.

## Acknowledgements

The authors would like thank The World Bee Project CIC and Hivetracks for providing funding for the purchase of the microscope(s).

## References

1. Bishop, S.P., Drummond, J.L.: Surface morphology and cell size measurement of isolated rat cardiac myocytes. Journal of molecular and cellular cardiology 11(5), 423–433 (1979)

2. Ineson, P.R.: Introduction to Practical Ore Microscopy. Routledge, ??? (2014)

3. Smith, C.E.: Pollen characteristics of african species of vernonia. Journal of the Arnold Arboretum 50(3), 469–477 (1969)

4. Walton, W., Beckett, S.: A microscope eyepiece graticule for the evaluation of fibrous dusts. The Annals of occupational hygiene 20(1), 19–23 (1977)

5. Verso, M.: The evolution of blood-counting techniques. Medical history 8(2), 149–158 (1964)

6. Lang, M., Rudolf, F., Stelling, J.: Use of youscope to implement systematic microscopy protocols. Current Protocols in Molecular Biology 98(1) (2012). doi:10.1002/0471142727.mb1421s98

7. Edelstein, A.D., Tsuchida, M.A., Amodaj, N., Pinkard, H., Vale, R.D., Stuurman, N.: Advanced methods of microscope control using *μ*manager software. Journal of Biological Methods 1(2), 10 (2014). doi:10.14440/jbm.2014.36

8. Koster, A., Chen, H., Sedat, J., Agard, D.: Automated microscopy for electron tomography. Ultramicroscopy 46(1-4), 207–227 (1992)

9. Ziese, U., Janssen, A., Murk, J.-L., Geerts, W., Van der Krift, T., Verkleij, A., Koster, A.: Automated high-throughput electron tomography by pre-calibration of image shifts. Journal of Microscopy 205(2), 187–200 (2002)

10. Boddeke, F., Van Vliet, L., Young, I.: Calibration of the automated z-axis of a microscope using focus functions. Journal of microscopy 186(3), 270–274 (1997)

11. He, P., Glowacki, G., Gkantiragas, A.: Unsupervised representations of pollen in bright-field microscopy. arXiv preprint arXiv:1908.01866 (2019)

12. He, P., Gkantiragas, A., Glowacki, G.: Honey authentication with machine learning augmented bright-field microscopy. arXiv preprint arXiv:1901.00516 (2018)

13. Mallat, S., Zhong, S.: Characterization of signals from multiscale edges. IEEE Transactions on Pattern Analysis & Machine Intelligence (7), 710–732 (1992)

14. Dedej, S., Hartfelder, K., Aumeier, P., Rosenkranz, P., Engels, W.: Caste determination is a sequential process: effect of larval age at grafting on ovariole number, hind leg size and cephalic volatiles in the honey bee (apis mellifera carnica). Journal of Apicultural Research 37(3), 183–190 (1998)

